# Label free fluorescence quantification of hydrolytic enzyme activity on native substrates reveal how lipase function depends on membrane curvature

**DOI:** 10.1101/2020.03.18.991711

**Authors:** Søren S.-R. Bohr, Camilla Thorlaksen, Ronja Marie Kühnel, Thomas Günther Pomorski, Nikos S. Hatzakis

## Abstract

Lipases are important hydrolytic enzymes used in a spectrum of technological applications, such as the pharmaceutical and detergent industry. Due to their versatile nature and ability to accept a broad range of substrates they have been extensively used for biotechnological and industrial applications. Current assays to measure lipase activity primarily rely on low sensitivity measurement of pH variations or visible changes on material properties, like hydration, and often require high amount of proteins. Fluorescent readouts on the other hand offer high contrast and even single molecule sensitivity, albeit they are reliant on fluorogenic substrates that structurally resemble the native ones. Here we present a method that combines the highly sensitive readout of fluorescent techniques while reporting enzymatic lipase function on native substrates. The method relies on embedding the environmentally sensitive fluorescent dye pHrodo and native substrates into the bilayer of liposomes. The charged products of the enzymatic hydrolysis alter the local membrane environment and thus the fluorescence intensity of pHrodo. The fluorescence can be accurately quantified and directly assigned to product formation and thus enzymatic activity. We illustrated the capacity of the assay to report function of diverse lipases and phospholipases both in a microplate setup and at the single particle level on individual nanoscale liposomes using Total Internal Reflection Fluorescence (TIRF). The parallelized sensitive readout of microscopy combined with the inherent polydispersity in sizes of liposomes allowed us to screen the effect of membrane curvature on lipase function and identify how mutations in the lid region control the membrane curvature dependent activity. We anticipate this methodology to be applicable for sensitive activity readouts for a spectrum of enzymes where the product of enzymatic reaction is charged.

## Introduction

Lipases are lipolytic enzymes, which catalyze the hydrolysis of ester bonds in medium to long triglycerides^1^ and have been studied vastly in the past^2–6^ for their broad range of applications from chiral organic synthesis^7^ to the production of bioethanol^8,9^, food production^10^ and cosmetics^11^. To support this plethora of functionalities lipases have been extensively engineered^12,13^, to meet the demand for increasingly specific chemical reactions^14^. Our capacity to optimize or alter protein function has radically improved through site directed mutagenesis of the active site^15,16^, directed evolution^17^, generation of novel functions using molecular dynamics (MD) simulations^18^ or machine learning^17^. The advancement of high throughput methodologies on the other hand may provide large numbers of mutants that have to be screened for the desired properties. Such screening creates the demand for quantitative and sensitive analysis of a desired function on native substrates without the need for protein labeling or substrate exchange.

Current understanding and evaluation of hydrolytic enzyme function relies primarily on ensemble assays^19^ like pH measurements^20^, simulations^6,21^ or model pre-fluorescent substrates in both solution^22–24^ and within thin films^24^. Ensemble methods report function on native substrates albeit they are insensitive and often require high amounts of substrates and protein^25,26^. Single particle tracking studies on the other hand report the spatiotemporal localization of enzymes, providing unprecedented insights on the behavior of diverse mutants but to date rely on correlations of diffusion to function^27–30^. Assays based on fluorogenic substrates offers low consumption and high sensitivity, even at the fundamental limit of single turnover cycles^31–33^, albeit at the cost of using substrate analogues that resemble the native enzyme substrates.

Here we present an assay that reports label free enzyme activity on native substrate systems while offering the high sensitivity of fluorescence readouts. The assay relies on implementation of the rhodamine-based pH sensitive fluorophore, pHrodo^34^, into liposome model membrane systems, together with lipase native substrates. Liposomes act as a 3D scaffold for substrate integration while offering an interface for the insertion of lipase amphipathic helix and interfacial activation^24,27–29,35,36^ even at relatively low concentration. Upon enzymatic ester hydrolysis, the produced charged carboxylic acids alter the local pH and charges at the membrane surface and consequently the fluorescent properties of the pHrodo, that can be quantified reproducibly using fluorescent techniques.

We firstly tested the utility of the assay for label free detection of enzyme activity using three different esterases Thermomyces Lanuginosus Lipase (TLL), TLL phospholipase A1 (TLL PLA1) and phospholipase A2 from honeybee venom (PLA2 bv) on their native substrates. We expanded the assay to single particle readout capitalizing on array of surface tethered vesicles^37,38^ and compared the native enzyme directly to both S146A TLL mutant and a mutant with the amphipathic helix in a locked and closed position (shut-up). The inherent polydispersity of the liposome’s sizes^32,37,38^ allowed us to evaluate the effect of liposome curvature on lipase function within a single experiment using nano moles of material. Our findings revealed S146A and shut-up contracts to display increased overall activity towards larger flatter vesicles, while the native enzyme showed no preference. The presented method allows for a sensitive and quantitative analysis platform for new lipase candidates on native substrate systems, offering in advance advanced mechanistic insights on protein recruitment and substrate dimensional preference.

## Results and discussion

To report the activity of hydrolytic esterases, such as TLL, we employed phospholipid liposomes as a model membrane system, previously shown to allow both binding and interfacial activation^24,29,32,35,39,40^ of lipases (see Fig. 1). The adaptability of the liposome setup enables facile variations in lipid composition and the incorporation of the pH-sensitive fluorescent dye pHrodo Red via a lipid anchor. The sensor pHrodo Red was selected as it has been employed to report local pH and charge variations^34,41,42^ on liposomes. The lipid-conjugated pHrodo Red is weakly fluorescent at pH 7.5, and as fatty acids are introduced by hydrolase activity we hypothesized that this would affect the local pH and charges resulting in fluorescent changes of the pHrodo sensor^34^ (see Fig. 1).

**Figure 1.**
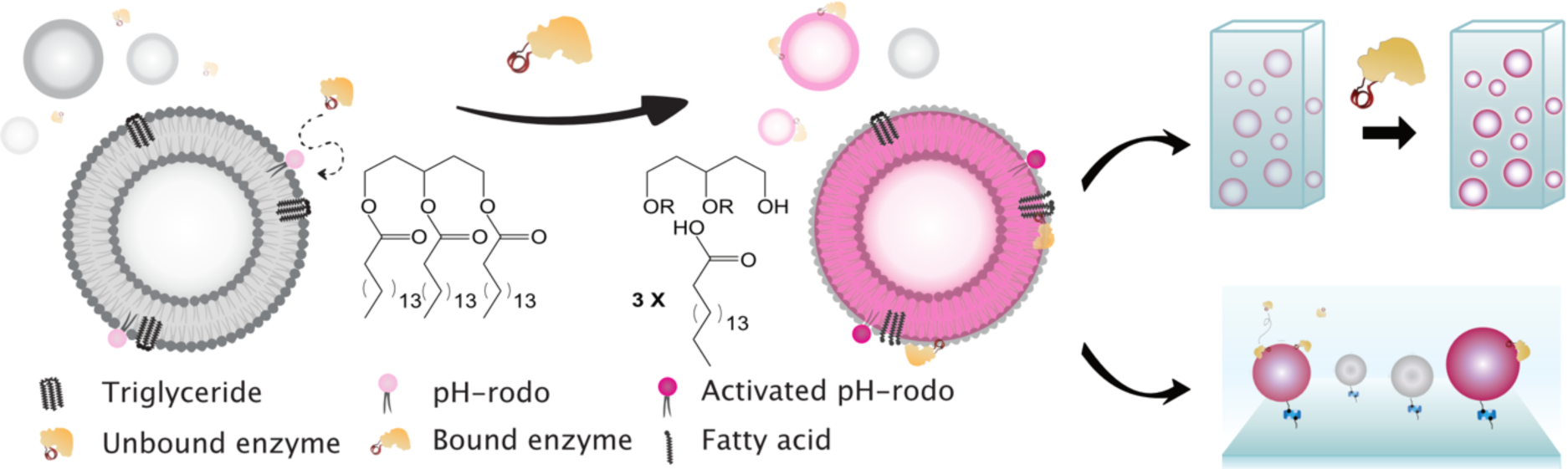
Imaging lipase activity utilizing model membranes (liposomes) containing the lipid-conjugated pH-sensitive fluorophore pHrodo Red and lipase substrates (triglycerides). Triglyceride hydrolysis by the lipase into glycerol and fatty acids changes locally the membrane environment affecting fluorescent properties of the pH sensor. Lipase activity are thereby reported in a label free manner. Using liposomes allow both ensemble measurements and single vesicle analysis by TIRFM.

### Recordings of conversion of triglycerides to fatty acids by TLL

We tested the utility of the assay by measuring the effect of TLL lipase addition to diester DOPC liposomes containing DOPS (8 mol%), the lipid-conjugated pH-sensitive fluorophore pHrodo Red (1 mol%) and the lipase substrate triolein (2 mol%) (Type 1 liposomes, see Table S1). Upon TLL addition, an increase in the fluorescence intensity of the pHrodo sensor was observed (Fig. 2A, black trace). To quantify the oleic acid formation, we calibrated the response of the pHrodo sensor to fatty acids presence in liposomes by generating liposomes containing increasing amounts of fatty acids and quantifying the pHrodo response (see Fig. S1). This methodology allowed us to quantify the time dependent production of oleic acid by lipase function using fluorescence readouts.

**Figure 2.**
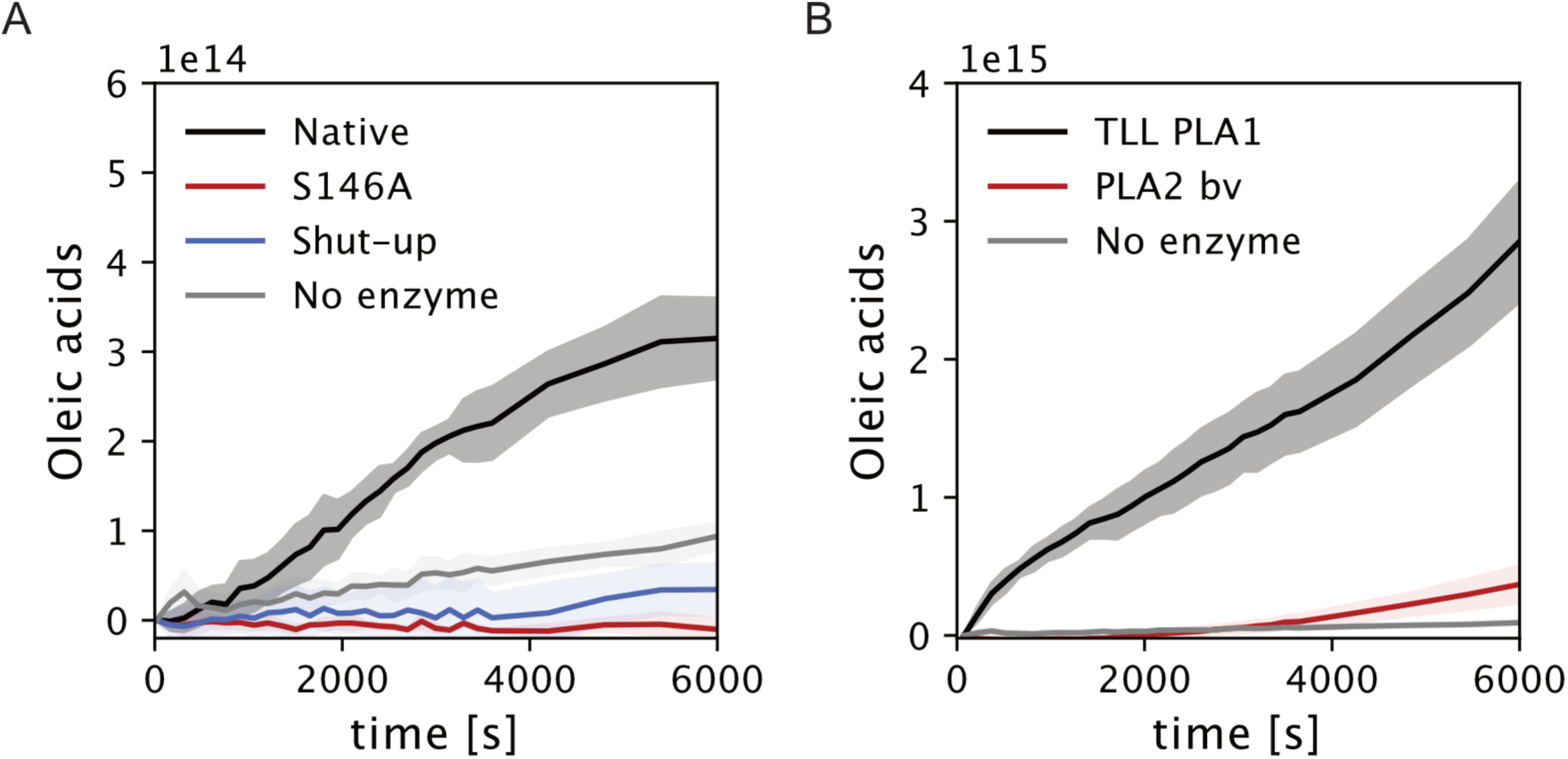
Assay validation and activity quantification for lipase variants as well as phospholipases on microplate reader setup. A) Time dependent oleic acid production by enzymatic hydrolysis of triolein integrated in liposomes by native TLLs (black line) (see Fig. S1 for calibration of intensity to oleic acids). Measurements of S146A mutant (red), Closed-lid (Shut-up) TLL C86C255 (blue) or no lipase (gray) display minimal differences on Type 1 liposomes, See Fig. S2 for Type 2. B) Expansion of the assay to report phospholipase activity. Both TLL PLA1 (black) and phospholipase A2 from bee venom PLA2 bv (red) produce a high number of oleic acids. Interestingly PLA2 bv displays similar activity rate to native TLL, while TLL PLA1 displays significantly higher activity in otherwise identical conditions. Control experiments performed without enzyme. Note the different y axis scale. Data from 3 independent measurements, shaded area corresponds to one standard deviation.

Several control experiments confirmed that the reported behavior indeed originate from lipase function. We initially tested the effect the S146A lipase variant that is structurally identical to native TLL contains a single mutation to the active serine and have been shown to bind similarly to native TLL on lipid interfaces^23,43,44^ but show minute activity^45^ (Fig. 2A, red trace). We found zero response which was similar to the blank control experiments with no enzyme addition (buffer alone) shown with grey trace in Fig. 2A. Lipases docking on liposomes does not affect the pHrodo response and is not biasing the recorded readout. We confirmed that the recorded response primarily originates from triolein hydrolysis by measurements on liposomes with non-hydrolysable ether lipids (Type 2 liposomes) (See Fig. S2). We also employed a closed lid TLL (shut-up) mutant where the amphipathic helix is locked in the closed conformation by a cysteine bridge ^40^. Similarly, no measurable response was observed for this mutant confirming the utility of the assay to record lipase function on native substrates (Fig 2A, blue trace).

### Method adaptability to report the functionality of diverse ester hydrolyzing enzymes

We then examined the assay adaptability to record the function of additional hydrolytic enzymes such as phospholipases. Consistent with our hypothesis our tests on phospholipase A_2_ from honeybee venom (PLA2 bv), revealed an enzyme dependent response from oleic acid production on Type 1 liposomes (Fig. 2B, red trace). The observed activity is similar to the activity of TLL under otherwise identical conditions. This is interesting as PLA2 bv may hydrolyze the phospholipids (DOPC and DOPS) of liposomes that are in large excess and as such has increased substrate available as compared to TLL, that only hydrolyzes the embedded triolein. TLL PLA1 variant^28^, known to both hydrolyze phospholipids and triglycerides (see Fig. 2B), displayed a ∼10-fold increase in oleic acid production, compared to native TLL and PLA2 bv. The fact that the assay reliably reports the activity of diverse hydrolytic enzymes (lipases and phospholipases) under several conditions, confirms its utility as a functional screening tool for new variants.

### Lipase activity recordings on single liposomes

We employed TIRFM to image in a parallel manner lipase function on individual liposomes. We tethered liposomes on passivated glass slides using biotin interactions (see Fig. 3A)^38^. This methodology maintains the spherical shape and structural integrity of liposomes under immobilization conditions^37,46^ and allows the unhindered binding of protein and protein motifs^32,37^. The method’s sensitivity^32,38,47^ enables mechanistic analysis of liposome curvature and its effect on lipase function^37^. While liposomes have sub-resolution dimensions, integration of pHrodo Red intensity prior to addition of lipase allows the accurate quantification of their nanoscale sizes^37^. Continuously integrating pHrodo intensity over the experimental time frame (for 16 min with a temporal resolution of 0.167 Hz) allows recording of the time dependent change in intensity that correlates with the lipase function on each liposome. Extraction of liposome intensity was done using a methodology we developed earlier^38^. Briefly, images were corrected for potential drift using a single particle tracking approach, after which the background corrected intensity is extracted (see experimental). Enzymatic activity was triggered by addition of enzyme at t= 2 min (see Fig. 3C, arrow). To ensure system saturation all concentrations of enzyme was significantly above the K_d_ for amphipathic helix docking on liposomes, as we have shown earlier^37^.

**Figure 3.**
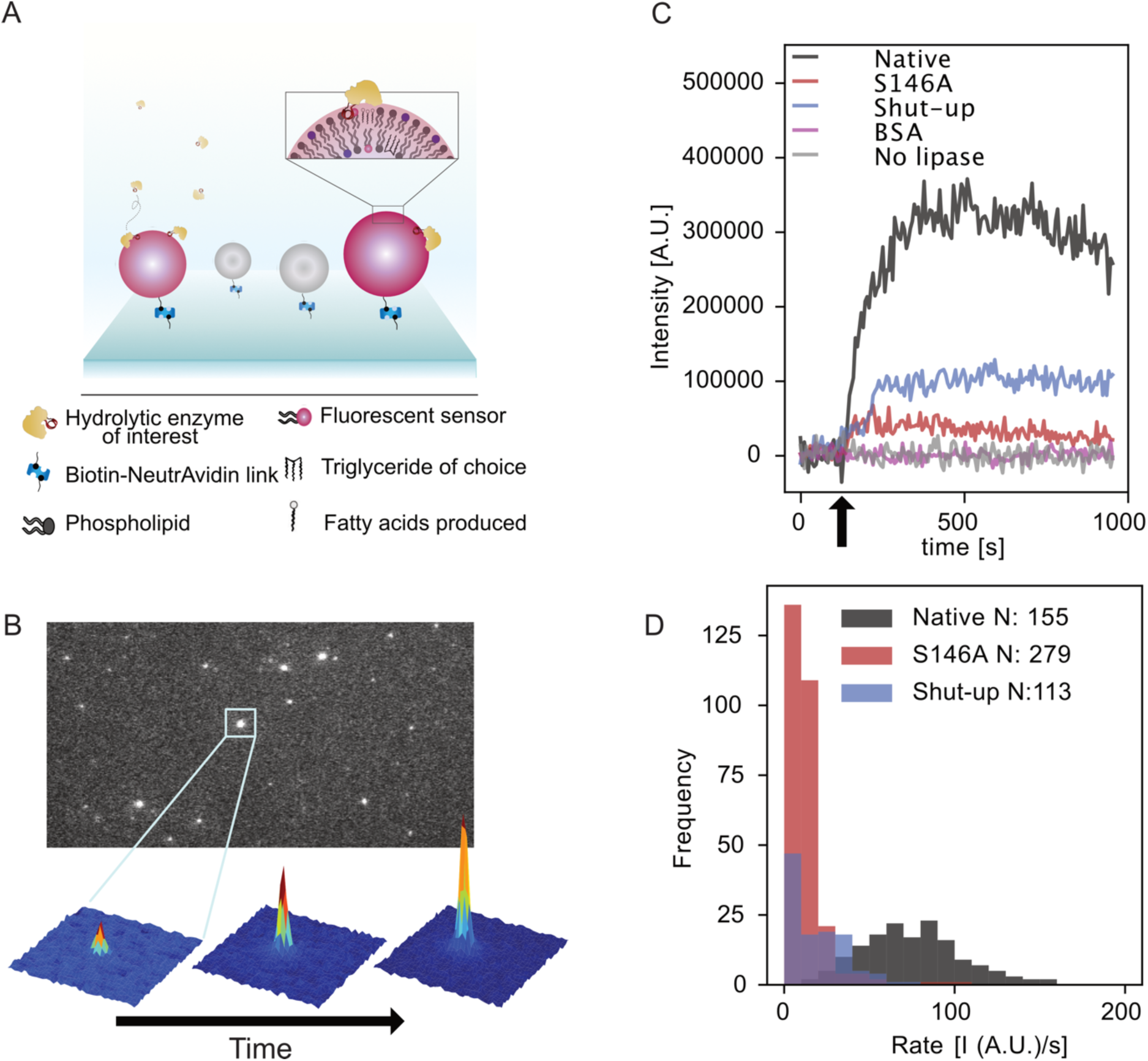
Parallel imaging of lipase activity on individual surface-tethered liposomes using TIRFM. A) Illustration of liposomes containing lipid-conjugated pHrodo and a lipase substrate (triglycerides) tethered to a passivated glass surface by NeutrAvidin. Zoom: Binding of lipase and enzymatic hydrolysis introduces fatty acids and charges that alter the local environment, thus resulting in a fluorescent response by the pH sensor. B) Representative TIRFM image of a surface with the immobilized liposomes. pHrodo implementation serves to localize and extract the dimension of sub-resolution particles. Zoom-in on a single liposome showing an intensity increase, due to product formation, as a function of time after lipase addition. C) Representative traces of liposomes with either addition of native TLL (black), S146A TLL (red), shut-up TLL (blue), buffer (no lipase, gray) or BSA (magenta) all normalized to initial intensity. Arrow indicate time of addition (see Fig. S3 for additional traces). Fitting active traces with an exponential function from t = 2 min until experiment end or liposome dissociation allowed relative rate extraction at each individual liposome (see experimental and Fig. S4). D) Distribution of extracted rates for lipase variants.

Enzyme additions resulted in an increase in pHrodo fluorescence intensity as displayed in the raw microscopy data zoom in Fig. 3B. We found ∼23-60% of liposomes to display intensity response, indicating a low copy number of enzymes were docked on each of the liposome. The intensity traces of liposomes not displaying lipase activity and control liposomes without enzyme illustrate practically zero effect of bleaching within the experimental time frame (see Fig. S5). Representative traces displaying the reliable recordings of enzymatic activity on individual liposomes by time dependent intensity increase are displayed in Fig. 3C.

Control experiments without enzyme revealed no detectable intensity increase, see Table 1 and Fig. 3C, compared to ∼30% of vesicles in experiments using native TLL. Additionally, the non-catalytic protein BSA, earlier shown to bind to liposomes^48^, resulted in no fluorescent increase (see Fig. S6). Addition of S146A and shut-up variant resulted in a minute increase in fluorescent intensity (see Fig. S3 for additional traces). Consistent with lipase activity we found ∼85% of liposomes to be disrupted after prolonged exposure to lipases see Fig. S3) resulting in loss of signal from the microscope surface. In all cases, liposome disruption only occurs after signal saturation and hence does not affect our readout. This phenomenon was not visible for S146A and shut-up mutants confirming lipase activity to cause liposome disruption.

**Table 1.** Event frequency and rates for the lipase variant in the TIRFM assay. Results are the means ± SD of 4 experiments.

| Lipase | Total liposomes | Events | Rate [1/s] |
| --- | --- | --- | --- |
| Native | 519 | 155 | $75.69 \pm 28.85$ |
| Shut-up | 541 | 113 | $19.33 \pm 15.47$ |
| S146A | 487 | 279 | $12.60 \pm 12.18$ |
| BSA | 600 | 0 |  |
| No Lipase | 531 | 0 |  |

We then compare the rate of catalysis towards individual liposomes for multiple TLL constructs, which revealed them to display different activities. As shown in Fig. 3, the intensity increases exponentially upon enzymatic addition, in agreement with our ensemble measurements and our earlier activity studies on individual liposomes^32^. Fitting to an exponential function all liposomes displaying an increase allowed us to extract the relative lipase rate displayed on each liposome (see materials and methods and Fig. S4 for trace fitting. Inspection of rates collected from native TLL (see Fig. 3D) revealed a mean rate of 76.68 I/s, while the shut-up mutant displayed a rate ∼ 4 times lower than the native enzyme from 75.69 ± 28.85 I/s to 19.33 ± 15.47 I/s (see Table 1, Fig. 3C and D). Addition of the S146A mutant resulted in an even lower rate of 12.60 ± 12.18 I/s, ∼ 6 times lower than the native enzyme as expected for the practically inactive variants^40,45^ (see Fig. S7 for all individual histograms of rates). We note that the recorded intensity increase may be convolution of two phenomena which we cannot resolve directly: the lipase binding densities on each liposome, and the activity of each of the bound lipases. The fact that we observe three distinct rate distributions (see supplementary Table 4) confirming the sensitivity of the assay to report differentiable activities from multiple enzymes.

### Single liposome setup reveals how lipase activity depends on membrane curvature

The active site of lipases is covered by an amphipathic helix and curvature can induce altered binding densities^35,49^ for species containing such a motif^37^. Such phenomena may be obscured by ensemble techniques as liposome populations are polydisperse in size and we have shown the diameters of a population extruded at 400 nm can vary 10 fold, from 40 to 400nm^37,38,46^. The single particle measurements allowed us to observe directly intensity variation in each liposome and compare the effect of liposome size and thus membrane curvature between the three lipase variants tested here (see Fig. 4A). A basic prerequisite for this is the monodisperse distribution of incorporated triolein lipid independently of liposomes diameters that we have reported recently^46^. Our recordings revealed that out of the 2000 analyzed liposomes 547 displayed enzymatic activity (see Table 1), in agreement with earlier results that only a fraction of liposomes will display docking and activity^37,50^. Consistent with our spectrometer assays the native displayed significantly higher activity than the S146A and shut-up variants for all membrane curvatures.

**Fig 4:**
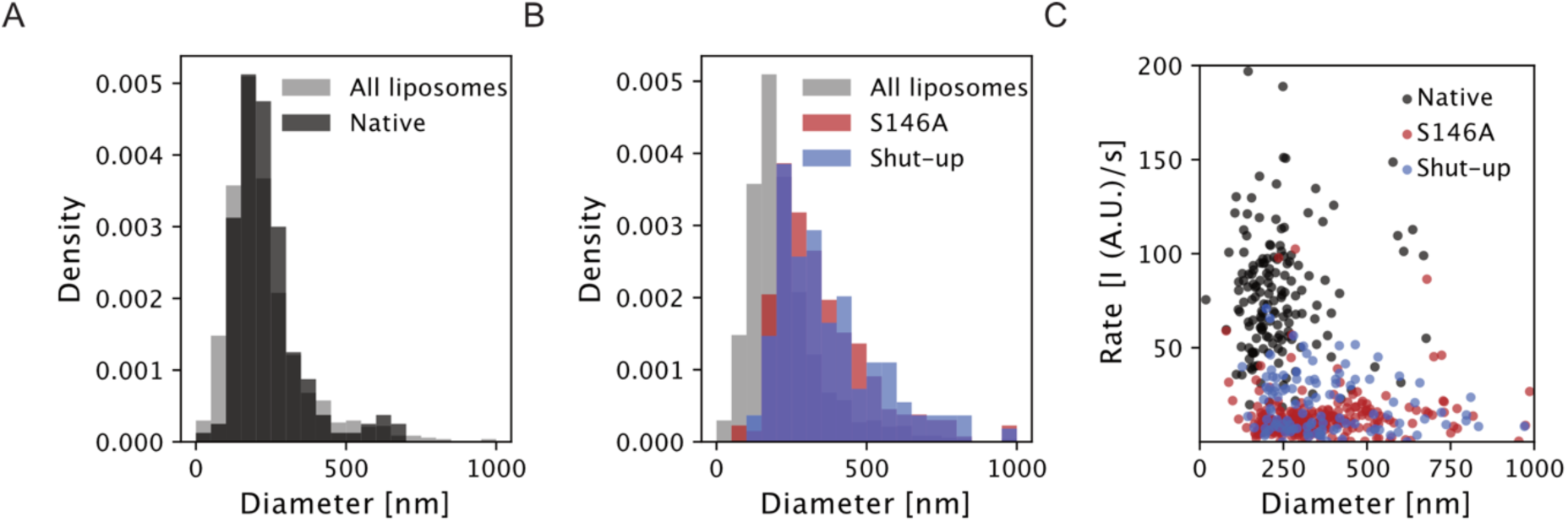
Lipase activity to size and curvature correlation A) Scatter plot of rate versus liposome size. B) Size distribution of total liposome population (gray) and liposomes displaying a signal from native TLL (black). Investigation reveals that the native enzyme will choose randomly from the available distribution. C) Size distributions for shut-up variant (blue) and S146A lipase (red) display both phenotypes to prefer larger vesicles, as compared to the entire population (grey).

Direct comparison of the original liposome population and the subpopulation where the native TLL displayed activity, shows no significant preference for size (see Fig. 4A, supporting table 2 and Fig. S8), as distributions are indiscernible. This is interesting as we and others have shown amphipathic helices, like the lid of TLL, to preferentially bind to areas of high curvature^37,51,52^. Plotting however the relative activity for each liposome size revealed minor dependence of activity to low membrane curvature for the native variant (Pearson correlation coefficient 0.012, p-value = 0.8). Analysis of the curvature preference of S146A lipase and shut-up lipase (see Fig. 4B) report these to be indistinguishable from each other. Both variants however are significantly different from both native and the parent population and display a shift towards higher sizes (see supplementary Table 2). One may argue this shift is due to the low activity of these variants, and that only larger vesicles contain the required high number of enzymes and substrate molecules to attain high, reliable signal. Careful inspection of Fig. 4C reveals that the S146A and shut-up lipase variants dock on all liposome sizes suggesting this is not the case. Interestingly examining the curvature dependent activity of the S146A variant (red) showed slight negative correlation (Pearson correlation coefficient - 0.065) suggesting the low activity of the S146A variant to be slightly increased towards smaller vesicles. No significant correlation of was observed for the shut-up variant (see supplementary Table 3 and Fig. S9).

The fact that only a fraction of (23-60%) of liposomes displayed docking^37,50^, indicates that while the presented assay cannot distinguish binding from activity, it is most likely reporting the signal from a low copy number of enzymes. Further optimization of the assay may offer the exciting possibility of recording the activity of single enzymes.

## Conclusion

Robust and label free quantification of activity for new protein constructs is crucial for developing catalysts with tailored functionalities. The advent of advanced engineering methodologies and even machine learning approaches^17^ have far outpaced the development of rapid, robust and sensitive methodologies to reliably analyze esterase behavior. Current methods are often either time demanding or insensitive and thus require high amounts of enzyme to make viable conclusions. Fluorescent technologies on the other hand are sensitive and require low enzyme amounts albeit require fluorogenic and thus often non-native substrates. The assay presented here address these potential shortcomings and offers a sensitive and label free lipase activity readout on native substrates. Liposome are ideally acting both as substrate scaffolds and as a model system for interfacial lipase activation^24,32,35,36,39^. Here we demonstrated the quantitative utility of this method to assay the activity of multiple lipase and phospholipases variants.

A significant advantage of the method is that it conveniently integrates sensitive quantitative microscopy with enzyme screening using parallelized imaging on arrays of surface tethered liposomes. The combination of membrane heterogeneities and protein concentration may create distinct combinatorial permutations of regulatory inputs, which can be screened while consuming only a few nano moles of protein. The single particle readout turns such inherent heterogeneities from a prohibitive problem in classical approaches to an experimental advantage in our assay allowing within a single experiment to screen the effect of membrane curvature on lipase function. Our method revealed that native TLL displayed no preference for liposome size but its activity was found to marginally dependent of liposome curvature. The observed activity on each liposome for the S146A and shut-up variant on the other hand, displayed a small but significant preference for larger liposomes.

Implementation of alternative enzyme substrates whose product locally alters charges on the membrane, or alternative membrane bound pH reporters emitting in different wavelengths may significantly expand the repertoire of the assay to account for a host of enzyme systems. The anticipated mechanistic insights attained in rapid and robust way could significantly speed up both characterization of future enzyme candidates and simultaneously aid the design of enzymes with custom functionalities designed to our needs. Expansion of the current assay could allow the parallelized recordings of both function but also molecular recruitment details for additional hydrolytic enzymes, whose substrate can be implemented within membrane surfaces.

## Experimental

### Materials

Trizma base (T1503; Sigma-Aldrich, Broendby, Denmark), Calcium chloride dihydrate (C3306; Sigma Aldrich), 1,2-di-O-(9Z-octadecenyl)-*sn*-glycero-3-phosphocholine (diether-DOPC; 999989; Avanti Polar lipids Inc., Alabaster, AL, USA), 1,2-dioleoyl-*sn*-glycero-3-phosphocholine (DOPC; 850375; Avanti Polar Lipids Inc.), 1,2-dioleoyl-*sn*-glycero-3-phospho-L-serine (DOPS; 840035C; Avanti Polar Lipids Inc.), 1,2-disteroyl-*sn*-glycero-3-phosphoethanolamine-N-[biotinyl(polyethylene glycol)-2000] (ammonium salt) (DSPE-PEG_2000_-biotin; Avanti Polar Lipids Inc.), Triolein (1,2,3-Tri(cis-9-octadecenoyl)glycerol; T7140; Sigma-Aldrich), Oleic acid (cis-9-Octadecenoic acid; O1008; Sigma Aldrich). Dioleoylphosphatidylethanolamine-conjugated pHrodo Red (DOPE-pHrodo Red) was synthesized as described earlier^41^.

### Enzyme variants

All lipase variants from Thermomyces Lanuginosus lipase (TLL) were kindly provided by Allan Svendsen (Novozymes A/S, Bagsvaerd): catalytic active TLL (D137C) with an aspartic acid to cysteine substitution allowing or potential labelling, practically catalytic Inactive TLL (S146A) with a serine to alanine substitution^43^, “Shut-Up” TLL with strategically mutated cysteine’s keeping the ‘lid’ locked in closed position restricting interfacial binding and access of substrate to the active site^40^, TLL PLA1 a TLL variant engineered to possess phospholipase activity^28^. Secretory phospholipase A_2_ from honey bee venom (P9279) was purchased from Sigma Aldrich.

### Liposome preparation

Liposomes were prepared by the thin-film hydration method followed by extrusion, as described earlier^32,37^. Briefly, lipids in chloroform were mixed in glass vial s to achieve the indicated lipid composition (Suppl. Table S1). A thin lipid film was formed by solvent evaporation for 10 min under constant nitrogen flow, followed by incubation under vacuum for at least 1 h. The thin lipid film was rehydrated with 50 mM Tris buffer adjusted to pH 7.5 with 1 M HCl for a final liposome concentration of 1 g/L. The vial was vortexed to ensure complete dissolution and left for 30 min at room temperature to allow liposome self-assembly. The resulting liposomes were passed 10 times through 50 nm size nucleopore polycarbonate membranes (Nuclepore™, 7026006, Whatman GmbH, Germany) supported by two filters on each side (610014; Avanti Polar Lipids Inc.) and mounted in a mini-extruder (610000, Avanti Polar Lipids, Alabaster, AL). Subsequently, the liposomes were freeze-thawed (cycle of 10 repetitions), as described earlier^37,38^ to ensure unilamellarity. Liposomes were flash-frozen in liquid nitrogen and kept at −20°C in a UV-protected box until measurements.

### Liposome composition

All liposomes were based on DOPC or non-hydrolyzable diether-DOPC (88 mol%; Suppl. Table S1). Additionally, the liposomes contained 1 mol% of the pH sensitive fluorescent probe DOPE-pHrodo Red for indirect reporting of lipase activity upon substrate hydrolysis. As lipase native substrate, 2 mol% triolein was incorporated into the liposomes. To study the effect of charged lipids on lipase association and activity, liposomes contained 8 mol% DOPS. To enable surface passivation of the liposomes for microscopy measurements, all liposomes contained 0.5 mol% DSPE-PEG-biotin.

### Ensemble lipase activity assay

The functionality of the assay was investigated on a range of liposome systems and enzyme variants in microtiter plates (black flat NUNC 384, 262260; Thermo Scientific). To each well, 50 µL Tris buffer (50 mM, pH 7.5) was added with 2.5 µL of liposomes (1 g/L) and 10 µL lipase variant (1.3 µM final concentration). For wells containing TLL PLA1 and PLA2 bv, 2 µL of 20 mM CaCl_2_·2xH_2_O) was added. Upon addition of enzyme, the plate was placed in the plate reader (Varioscan flash 3001, Thermo Scientific). The fluorescence intensity of DOPE-pHrodo Red was measured at 585/10 nm using an excitation wavelength of 532/10 nm every 2.5 min for the first hour, then every 10 min for 2 hours. All measurements were carried out at 25°C.

### Calibration assay

To directly correlate Relative Fluorescence Units (RFU) with the total amount of fatty acids produced liposomes containing an increasing amount (0.5, 1, 2, 4 and 6 mol%) of oleic acid were prepared as described above. Oleic acid presence may locally alter both pH and the charges of the system. Calibrating the observed response to the amount of oleic acid calibrates the overall response of the liposome system to product formations. Re-hydration with Tris buffer (50 mM, pH 7.5) occurred just prior to measuring to ensure quantitative fatty acid partitioning into the membrane^37,53,54^. Experiments were performed within 30min after vesicle preparation to ensure that the slow^55^ desorption of oleic acids from vesicles does not impair the calibration curves. Due to the longer time scale requirement for microscopy measurements, for surface immobilization and washing and microscope setting, only relative rates are recorded. The linearity of the calibration curves ensures that the relative intensity variation recorded for different variants correspond directly to product rate variations.

### Single liposome fluorescence microscopy

For fluorescence microscopy, a flow chamber was assembled. Glass coverslips (No. 1.5, ibidi GmbH) were activated by plasma edging in vacuum, attached to the bottom of a microfluidic chamber (channel width, 3.8 mm; channel height, 0.4 mm; sticky-Slide VI 0.4, ibidi GmbH) and sequentially passivated as described earlier^38,56^. The chamber was flushed with 1 mL buffer followed by the liposome solution. Liposomes were diluted to achieve a density of around 100 liposomes pr. field-of-view and sequentially washed to remove non-bound liposomes. The fluorescence microscopy was performed using an Olympus (IX 83, Olympus) Total Internal Reflection Fluorescence (TIRF) microscope with a Hamamatsu (ImagEM X2, Hamamatsu) EMCCD camera and a 100x objective/N.A. 1.49 (UAPON 100XOTIRF, Olympus), resulting in a pixel size of 160 nm. Images were recorded every six seconds for 20 min using a solid-state laser line (excitation 532 nm) and 80 ms exposure time. The penetration depth was set to 80 nm to ensure excitation of only liposomes and thus reduce background noise. At t=2 min either buffer alone (control) or enzyme (47 μM TLL, 50 μM shut-up, 8 μM S146A, or 5 µM BSA) was flowed into the chamber; while continuous recording. To achieve a higher throughput, four individual spots were imaged sequentially using the motorized stage.

### Extracting intensities from single liposome assay

Extraction of fluorescent intensity traces from individual liposomes where done using in-house developed routines in python, following recently published methodologies^38,57^. Briefly, time series were firstly corrected for potential drift from addition of enzyme as recently published^38^. After ensuring that no drift was observed, the full trajectories would be collected and individually corrected for potential background contributions as also shown in^38^, resulting in trajectories as displayed in Fig. 3 and Fig. S3. Conversion of liposome intensity to absolute size were done as described by us earlier^37,38^, using a correlation factor, F, of 0.9611 found using Dynamic Light Scattering (DLS).

### Fitting activity traces from single liposome assay

To quantify the enzymatic activity on liposomes, traces displaying an increase (see Fig. 3) were fitted to the following equation:

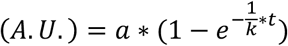

where a is a scaling factor, k is the rate decrease per second and t is the time. Each trace where fitted from lipase addition (t = 2 min) and until either the end of the experiment (t = 16 min), or the dissociation of the liposomes (see Fig. S4. All fitting was done using custom made routines in python.

## Acknowledgements

This work was funded by the Villum foundation via the young investigator fellowship (grant 10099) and the Bionec Center of excellence (grant 18333), the Carlsberg foundation Distinguished Associate professor program (CF16–0797) for NSH) Work at The Novo Nordisk Foundation Center for Protein Research (CPR) is funded by a generous donation from the Novo Nordisk Foundation (Grant number NNF14CC0001) T.P acknowledges a lab exchange grant from the Ruhr University Bochum (R.M.K).

## Author contributions

*SSRB and CT contributed equally. N.S.H designed and supervised the project. R.M.K synthesized DOPE-pHrodo Red. CT performed microplate reader and microscope measurements. CT and NSH designed the assay. SSRB performed microscopy measurements and analyzed data. SSRB with NSH and CT wrote the paper with inputs from all authors.

