## Supplementary information for "Label free fluorescence quantification of hydrolytic enzyme activity on native substrates reveal how lipase function depends on membrane curvature"

[a]Department of Chemistry & Nanoscience Center, University of Copenhagen, Thorvaldsensvej 40, Frederiksberg C 1871, Denmark. [b] Novo Nordisk Center for Protein Research (CPR), University of Copenhagen, Blegdamsvej 3B, Copenhagen 2200, Denmark. [c] Biophysics, Novo Nordisk A/S, Novo Nordisk Park 1, Maaloev 2760, Denmark. [d] Drug Delivery and Biophysics of Biopharmaceuticals, Department of Pharmacy, University of Copenhagen, Universitetsparken 2, Copenhagen 2100, Denmark. [e] Faculty of Chemistry and Biochemistry, Department of Molecular Biochemistry, Ruhr University Bochum, Universitätstrasse 150, D-44780 Bochum, Germany. [f] Department of Plant and Environmental Sciences, University of Copenhagen, Thorvaldsensvej 40, 1871 Frederiksberg C, Denmark

Supplementary Table S1. Liposome species used. All preparations additionally include 1 mol% DOPE-pHrodo Red and 0.5 mol% DSPE-$PEG_{2000}$-biotin.

| Liposome type | Triglyceride (2 mol%) | DOPS (8 mol%) | DOPC (88 mol%) |
|---|---|---|---|
| Type 1 | Triolein | diester | diester |
| Type 2 | Triolein | diester | diether |

Supplementary Figure S1. Converting intensity to oleic acid.

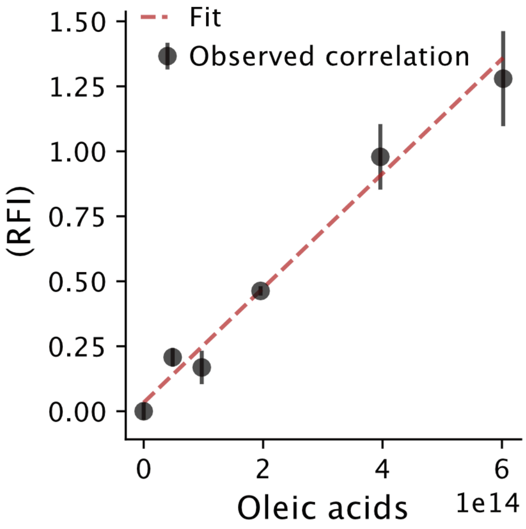


Supplementary Figure S1. Oleic acid calibration curve. Liposomes (type 2 containing increasing amounts of oleic acid together with 1 mol% DOPE-pHrodo Red, 8 mol% DOPS, 0.5 mol% DSPE-PEG$_{2000}$-biotin and diether DOPC. For each liposome preparation the corresponding fluorescence intensity at 585 nm was measured upon excitation at 532 nm. The relative fluorescence intensity (RFI) was normalized to zero by subtraction of the fluorescence intensity from liposomes prepared without oleic acids, but containing 2 mol% Triolein (which is the start condition for all of the kinetic measurements). Due to fatty acid dissociating from liposomes, the calibration could not be performed for the single liposome setup. Error bars denote standard deviation of 5 wells. All experiments were conducted in Tris buffer (50 mM, pH 7.5). The data was fitted with a linear function: RFI = 2.204·10-15 · OA + 0.0333, having chi-square = 6.64 and p = 0.23, revealing a linear correlation between fluorescent increase and the amount of fatty acids produced.

Supplementary Figure S2. Functionality towards different liposome systems

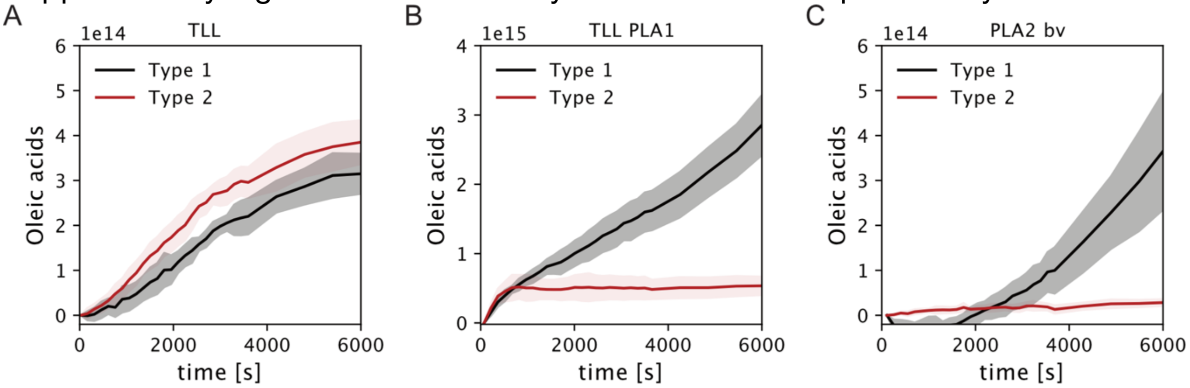


Supplementary Figure S2. Assay functionality on multiple liposome systems (see Table S1) measured by microplate measurements. A) Native TLL activity measured under different liposome compositions reveals no significant change from normal conditions (black) upon exchange of DOPC lipids with non-hydrolysable ether lipids (red). B) TLL phospholipase A1 (TLL PLA1) activity is dramatically reduced, as expected, upon removal of the bulk substrate DOPC (red) as compared to normal conditions (black). C) Phospholipase A2 from bee venom (PLA2 bv) activity likewise yielded diminished activity readouts from removal of DOPC substrate (red).

Supplementary Figure S3. Additional activity traces from single vesicles

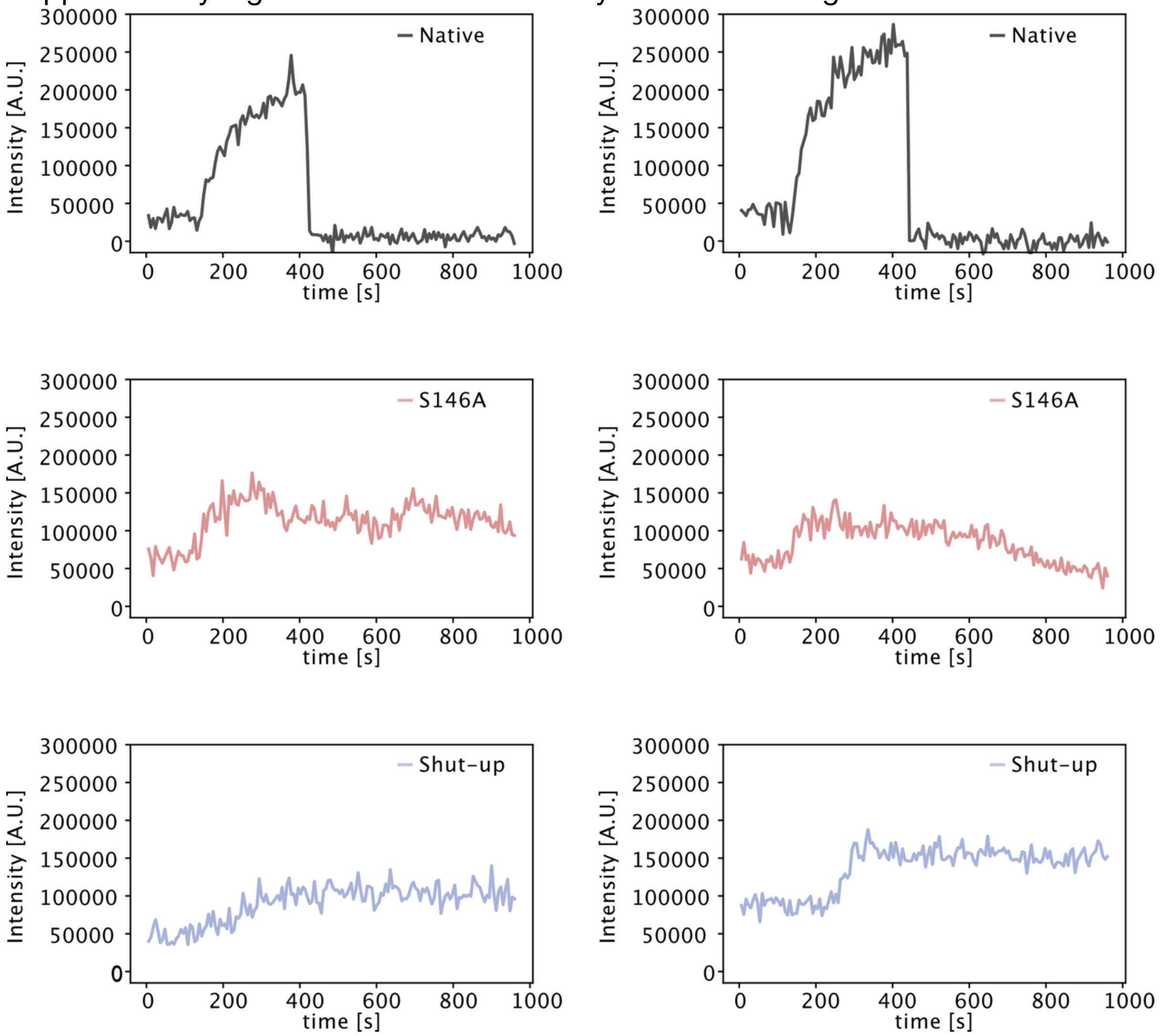


Supplementary Figure S3. Additional traces from single liposome assay. Top) Native TLL traces displaying that some liposomes display a dissociation after recorded enzymatic activity (~ 85 % of traces). Middle) S146A. Bottom) Shut-up.

Supplementary Figure S4. Rate fitting examples.

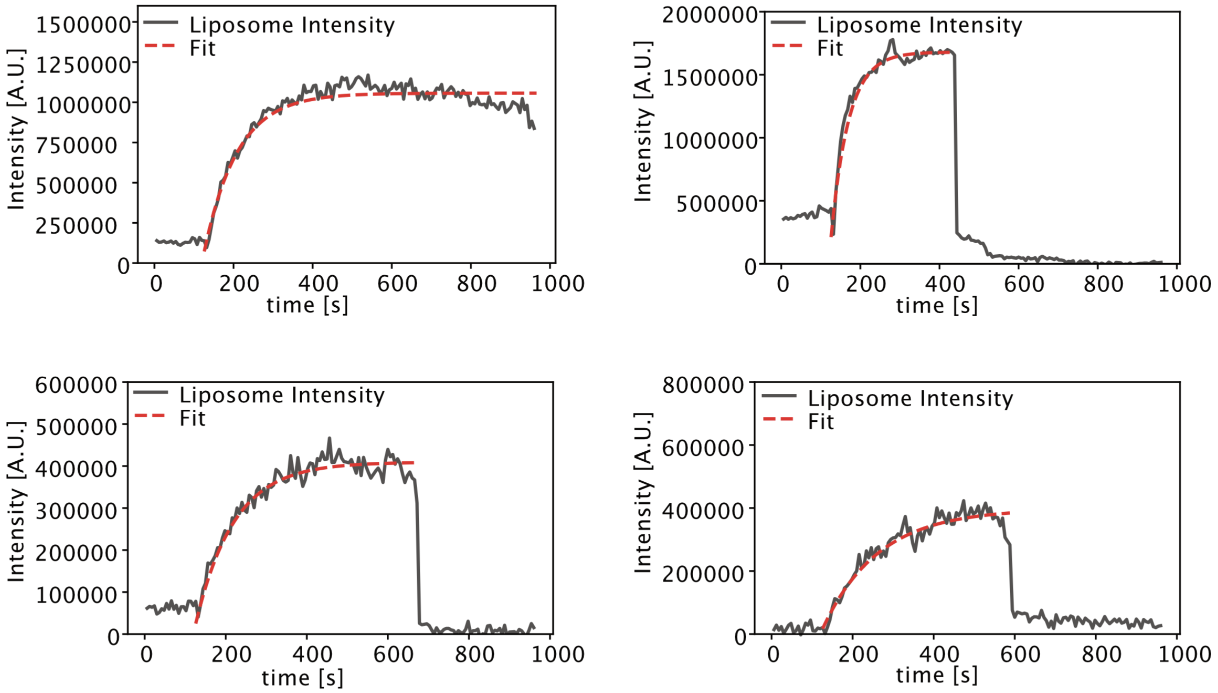


Supplementary Figure S4. Fitting liposome activity traces to exponential increase (see experimental section for detailed description and formula).

Supplementary Figure S5. Liposome stability.

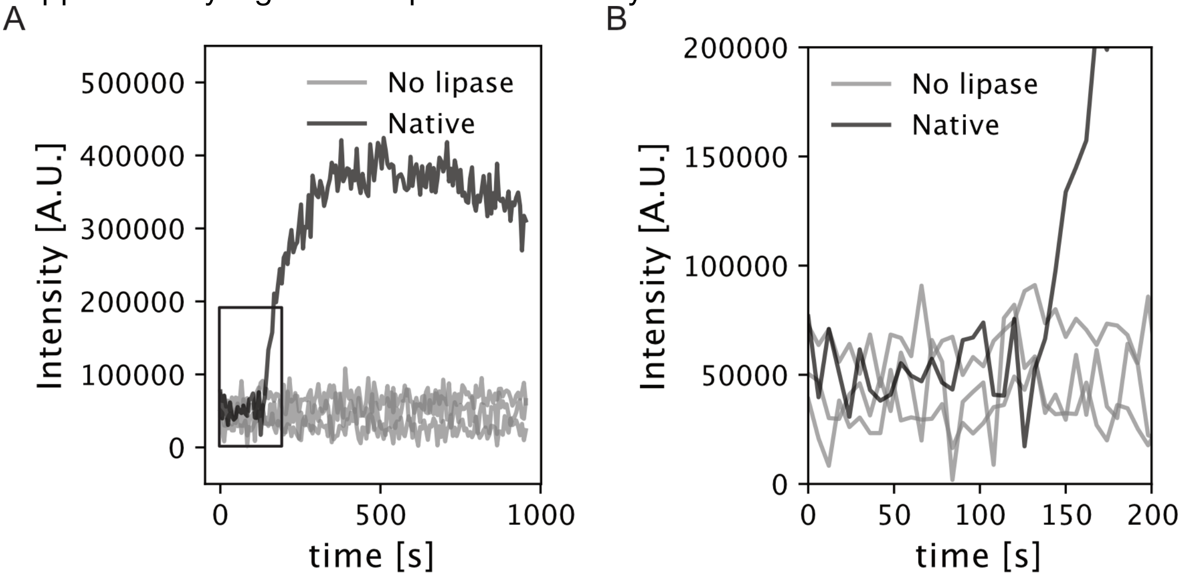


Supplementary Figure S5. Representative trajectories from single liposomes. A) Comparison between a liposome displaying an increase due to native lipase (black trace) and liposomes without lipase addition (buffer only, gray traces) reveals that bleaching is not biasing our data and liposomes are stable over the experimental time frame if no enzyme is added. B) Zoom in on square region from A.

Supplementary Figure S6. Addition of BSA to single liposome setup

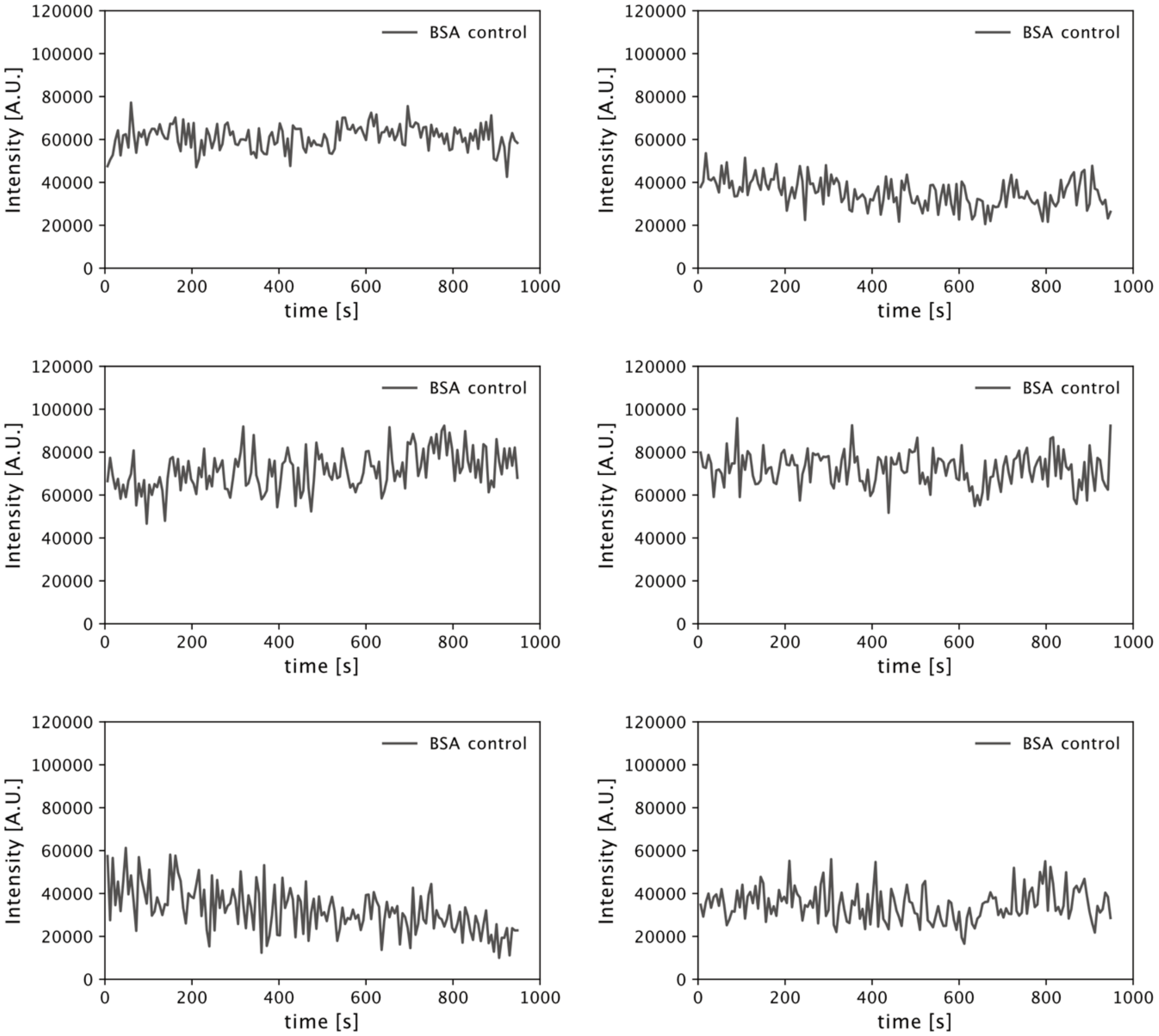


Supplementary Figure S6. Addition of BSA to single liposome setup. Representative traces from experiments with addition of BSA, displaying no increase in fluorescence, supporting the protein binding is not the sole underlying reason for intensity increase

Supplementary Figure S7. Distribution of relative activity rates for all conditions

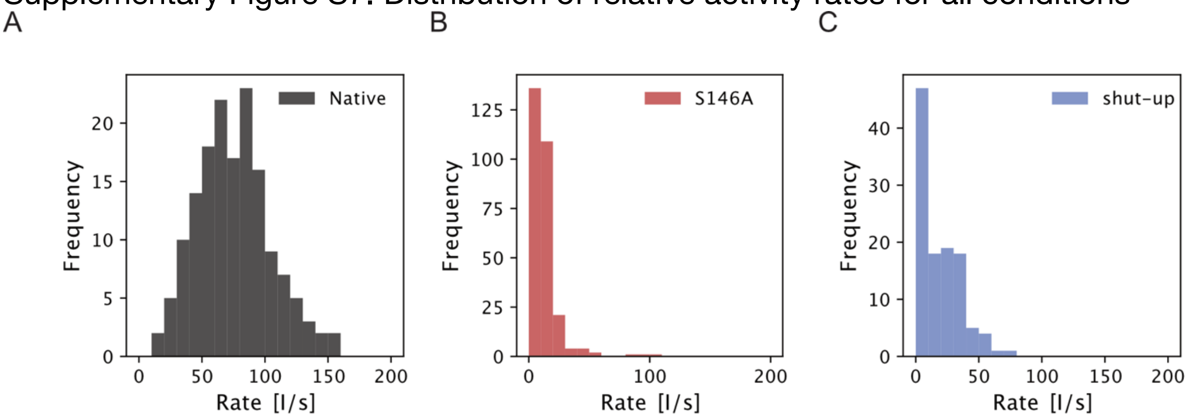


Supplementary Figure S7. Activity rate distributions for all experimental conditions. A) Native TLL. B) S146A TLL. C) Shut-up TLL.

Supplementary Figure S8. Liposome size distribution.

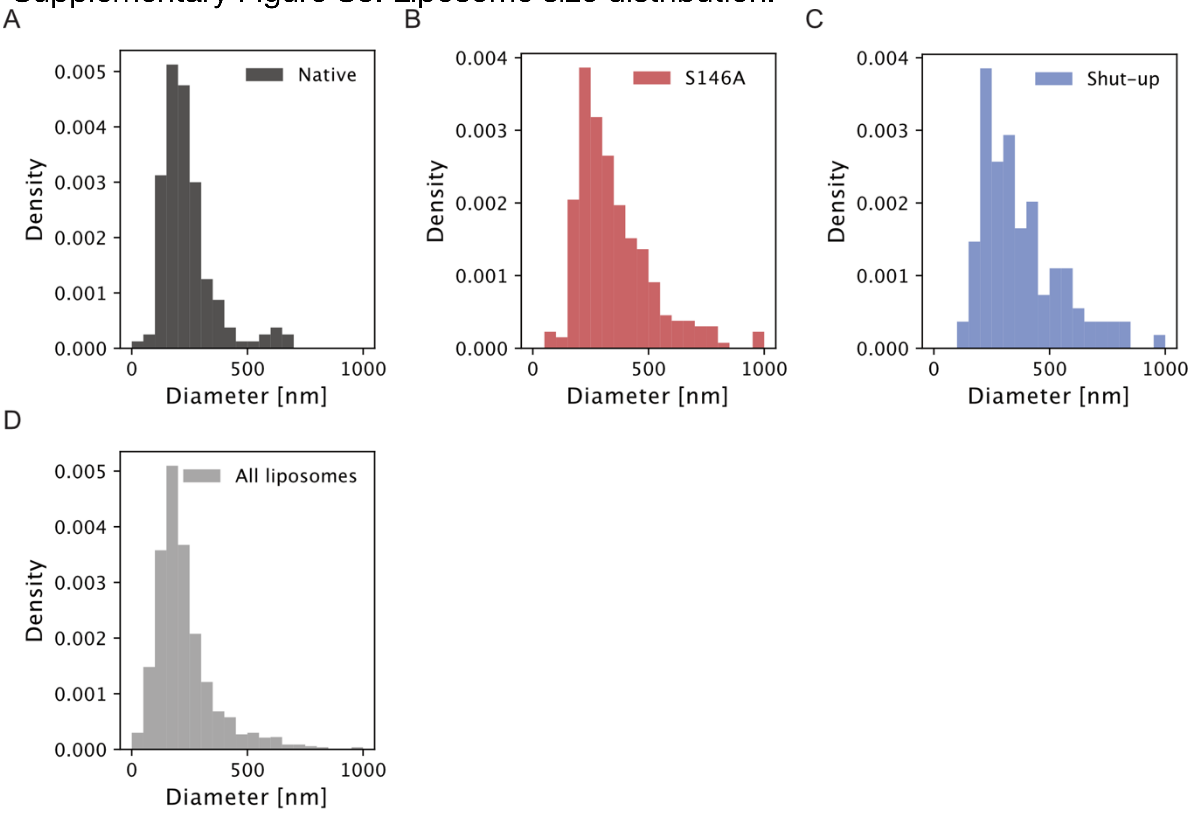


Supplementary Figure S8. Distribution of sizes for all liposomes recorded in the single liposome setup. A) Native. B) S146A. C) Shut-up. D) Parent liposome population (all liposomes measured).

Supplementary Table 2. Liposome size distribution comparison by student's t test.

| | All liposomes | All rates | Native | S146A | Shut-up |
|---|---|---|---|---|---|
| N | 2623* | 547** | 155** | 279** | 113** |
| All rates | <0.0001 | | | | |
| Native | 0.701 | <0.0001 | | | |
| S146A | <0.0001 | 0.0110 | <0.0001 | | |
| Shut-up | <0.0001 | 0.0560 | <0.0001 | 0.968 | |

* Data from all recorded liposomes

** Data from all liposomes displaying enzymatic activity

Supplementary Table 2. Size distribution statistics from single liposome measurements. All size distributions are compared to evaluate, which sub-distributions are from the same population.

Supplementary Figure S9. Liposome size to rate relation

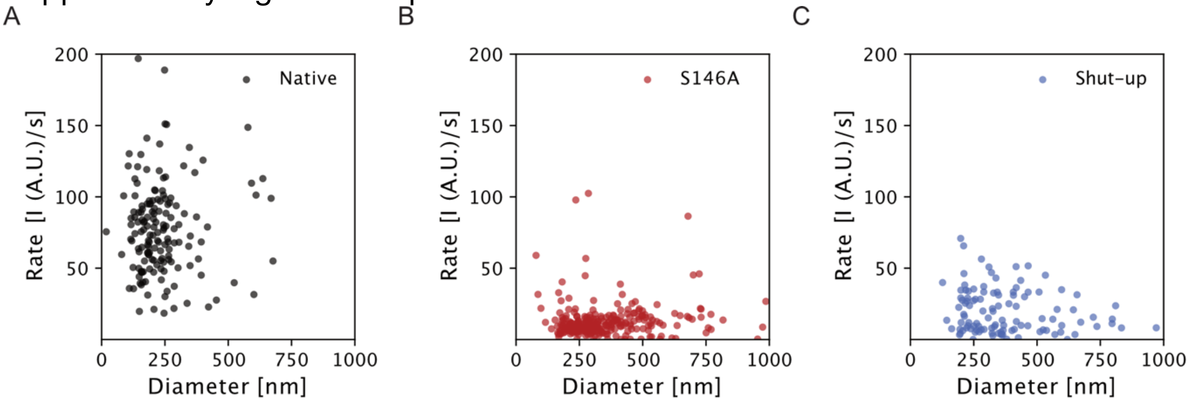


Supplementary Figure S9. Scatterplot displaying the relation between liposome size and observed activity rate. A) Native TLL. B) S146A TLL. C) Shut-up TLL. Pearson correlation coefficient and corresponding p-value are given in supplementary Table 3.

Supplementary Table. 3. Liposome size to rate relation by pearson correlation coefficient.

| | Pearson correlation coefficient | P-value |
|---|---|---|
| All rates | -0.256 | <0.0001 |
| Native | 0.012 | 0.803 |
| S146A | -0.065 | 0.274 |
| Shut-up | -0.253 | 0.007 |

Supplementary Table 3. Liposome size to activity rate correlation by pearson correlation coefficient.

Supplementary Table. 4. Comparison of activity rates by student's t-test.

| | All rates | Native | Inactive | Shut-up |
|---|---|---|---|---|
| N | 547 | 155 | 279 | 113 |
| Native | <0.0001 | | | |
| S146A | <0.0001 | <0.0001 | | |
| Shut-up | <0.0001 | <0.0001 | 0.013 | |
